# Quantification of the effect of site-specific histone acetylation on chromatin transcription rate

**DOI:** 10.1101/679944

**Authors:** Masatoshi Wakamori, Kohki Okabe, Kiyoe Ura, Takashi Funatsu, Masahiro Takinoue, Takashi Umehara

**Author notes:** To whom correspondence should be addressed. K.O., M.T. or T.U.

## Abstract

Eukaryotic transcription is epigenetically regulated by chromatin structure and post-translational modifications (PTMs). For example, lysine acetylation in histone H4 is correlated with activation of RNA polymerase I-, II-, and III-driven transcription from chromatin templates, which requires prior chromatin remodeling. However, quantitative understanding of the contribution of particular PTM states to the sequential steps of eukaryotic transcription has been hampered partially because reconstitution of a chromatin template with designed PTMs is difficult. In this study, we reconstituted a di-nucleosome with site-specifically acetylated or unmodified histone H4, which contained two copies of the *Xenopus* somatic 5S rRNA gene with addition of a unique sequence detectable by hybridization-assisted fluorescence correlation spectroscopy. Using a *Xenopus* oocyte nuclear extract, we analyzed the time course of accumulation of nascent 5S rRNA-derived transcripts generated on chromatin templates *in vitro.* Our mathematically described kinetic model and fitting analysis revealed that tetra-acetylation of histone H4 at K5/K8/K12/K16 increases the rate of transcriptionally competent chromatin formation ~3-fold in comparison with the absence of acetylation. We provide a kinetic model for quantitative evaluation of the contribution of epigenetic modifications to chromatin transcription.

## INTRODUCTION

Eukaryotic genes are transcribed by three classes of multisubunit DNA-dependent RNA polymerases (RNAPs) (1). Ribosomal RNA (rRNA) genes, protein-coding genes, and short non-coding genes, including those for 5S rRNA and transfer RNA (tRNA), are transcribed by RNAP I, II, and III, respectively (2–4). Because eukaryotic genomic DNA interacts with positively charged histone proteins and is compacted into chromatin, gene transcription is regulated not only by processes involving a naked DNA template—such as the recruitment of general transcription factors and RNAPs to the promoter, initiation, and elongation—but also by processes involving a chromatin template, such as chromatin accessibility for the transcription machinery and chromatin remodeling (5,6). The structural unit of chromatin, the nucleosome, is formed by wrapping DNA (145–147 bp) around the histone octamer, which consists of two copies of each of the core histones (H2A, H2B, H3, and H4) (7,8). Each core histone has a lysine (K)-rich N-terminal tail that protrudes through nucleosomal DNA, and these lysine residues are subjected to post-translational modifications (PTMs) such as acetylation, methylation, and ubiquitination (9). Lysine acetylation (Kac) in the histone H4 tail facilitates chromatin transcription by RNAPs I, II, and III in comparison with unmodified histone H4 (10–16); this may occur by enhancing the remodeling or repositioning (or both) of the nucleosome positioned near the transcription start site (17,18). Additionally, binding of an epigenetic reader protein, such as a bromodomain-containing protein, to chromatin may be facilitated by acetylation and may increase the rate of chromatin transcription (19,20).

The N-terminal tail of histone H4 has four major acetylation sites (K5, K8, K12, and K16), which are highly conserved from yeast to human. These lysines can be acetylated by histone acetyltransferases, and the number of acetylated residues correlates with RNA transcription activity (15,16,21). Among several H4 acetylation states (22–25), tetra-acetylation at K5/K8/K12/K16 is particularly important, because this hyperacetylated state is found in euchromatin regions where the nearby genes are transcriptionally most active (26). H4 acetylation at K5/K8/K12/K16 correlates with the expression of both the RNAP II- and III-transcribed genes (15,16). However, the contribution of each modification state of each histone to the sequential steps of chromatin transcription is yet to be quantified because of the difficulty in precise reconstitution of a chromatin template with the epigenetic modification(s) of interest (27). Using genetic code expansion and cell-free protein synthesis, we synthesized histone H4 containing designed site-specific acetylation(s) and reconstituted a H4-K5/K8/K12/K16-tetra-acetylated nucleosome (28,29). Despite the suggested importance of tetra-acetylation at K5/K8/K12/K16, this modification does not affect the crystal structure of the nucleosome core particle (29). Therefore, the effect of histone H4 acetylation on the dynamics of the nucleosome core needs to be analyzed.

Because the 5S rRNA gene can be chromatinized with a single nucleosome, it has been used as a model gene in an *in vitro* chromatin transcription assay (27,30). When whole histones including the H4 tail are acetylated in a di-nucleosome chromatin template containing a tandem of two 5S rRNA gene cassettes (*i.e.,* X5S 197-2 (27)), 5S rRNA transcription is activated *in vitro* (27). However, it is difficult to monitor transcription in real time because *in vitro* transcripts are usually detected by electrophoresis using radioisotope-labeled UTP.

In this study, we developed a di-nucleosome tandem *Xenopus* 5S rRNA gene cassette with a unique sequence derived from *c-fos.* Hybridization of end-initiated nascent transcripts containing two copies of this sequence with the corresponding antisense fluorescent probe enabled ultrasensitive real-time monitoring of transcript accumulation in a *Xenopus* oocyte nuclear extract by fluorescence correlation spectroscopy (FCS). We reconstituted the di-nucleosome chromatin template with non-acetylated or tetra-acetylated histone H4 and monitored accumulation of 5S transcripts. Our mathematically described kinetic model allowed us to determine the rates of chromatin transcription from non-acetylated and H4-tetra-acetylated chromatin templates. The methodology developed in this study can be used for quantitative analysis of the contribution of epigenetic modification(s) to chromatin transcription.

## MATERIALS AND METHODS

### Reconstitution of H4-acetylated histone octamers

Core histones were prepared and histone octamers were refolded essentially as described previously (31). Briefly, the full-length human histones H2A type 1-B/E, H2B type 1-J, and H3.1 were produced in *E. coli* BL21 (DE3) and purified by Ni-Sepharose affinity chromatography. K5/K8/K12/K16-acetylated histone H4 (hereafter referred as tetra-acetylated H4) was produced in an *E. coli* cell-free protein synthesis system with the expanded genetic code (28) as described (29). For refolding of histone octamers, equimolar amounts of histones (*i.e.,* H2A, H2B, H3, and unmodified or tetra-acetylated H4) were dissolved in 20 mM Tris-HCl buffer (pH 7.5) containing 6 M guanidine-HCl and 10 mM DTT, and dialyzed against 10 mM Tris-HCl buffer (pH 7.5) containing 2 M NaCl, 1 mM EDTA, and 5 mM 2-mercaptoethanol. The histone octamers were purified by size-exclusion chromatography on a Superdex 200 column (GE Healthcare).

### Preparation of di-nucleosome DNA

The 424-bp di-nucleosome DNA fragment X5S 197-2 composed of two tandem cassettes of the 197-bp *Xenopus borealis* somatic 5S rRNA gene with an upstream sequence (Figure 1A, top) was prepared as previously described (32). To produce the 479-bp di-nucleosome DNA fragment (X5S 217F-2; Figure 1A, bottom), two synthetic DNAs (250 bp and 229 bp) were purchased from Eurofins Genomics. The 250-bp DNA was the first 5S rRNA gene cassette (−108 to +142), in which a PvuII site (5’-CAGCT G-3’) was inserted at the 5’ end and the c-*fos* antisense probe sequence (5’-GCGGA GACAG ACCAA CTAGA-3’) was inserted at the +115 position. The 229-bp DNA was the second 5S rRNA gene cassette (+143 to +371), in which the c-*fos* antisense probe sequence was inserted at the +332 position and the PvuII site was inserted at the 3’ end. The two DNAs were ligated using a Gibson Assembly Master Mix (New England Biolabs) and subcloned into the pWMD01 vector (33). The di-nucleosome DNA with the modified 5S rRNA gene cassettes (X5S 217F-2) was excised with PvuII and purified by ion-exchange chromatography on a TSK-gel DEAE-5PW column (Tosoh Corporation). In the naked DNA transcription assay, the maxi 5S-2 template consisting of two tandem maxi 5S genes (27,34) was compared with X5S 197-2 and X5S 217F-2.

**Figure 1.**
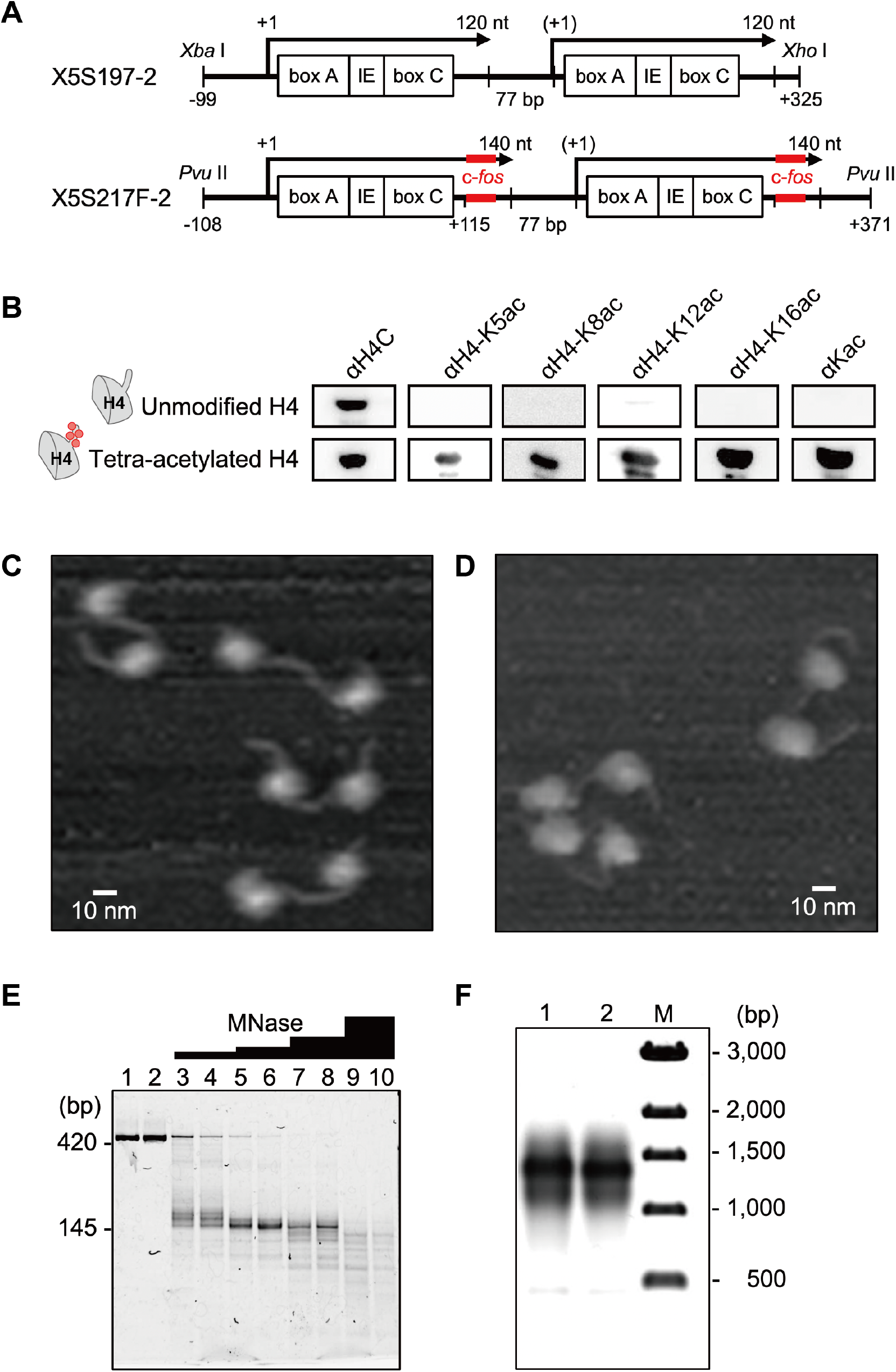
Reconstitution of H4-tetra-acetylated di-nucleosomes for chromatin transcription. (A) Scheme of di-nucleosome rRNA gene cassettes. Internal control regions (box A; IE, intermediate element; box C) of the 5S rRNA gene are indicated. The X5S 197-2 construct (27) (top) was modified by introducing a c-*fos* antisense probe sequence (bottom) for fluorescence correlation spectroscopy measurements. (B) Western blotting of unmodified and site-specifically tetra-acetylated histone H4 proteins. Kac, acetylated lysine. (C, D) Atomic force microscopy images of 5S rRNA di-nucleosomes reconstituted with unmodified histone H4 (C) or K5/K8/K12/K16-tetra-acetylated H4 (D). (E) Di-nucleosome digestion with micrococcal nuclease (MNase). Lanes 1, 3, 5, 7 and 9, di-nucleosomes with unmodified H4; lanes 2, 4, 6, 8 and 10, di-nucleosomes with K5/K8/K12/K16-tetra-acetylated H4. Lanes 1 and 2, di-nucleosomes were incubated in MNase reaction buffer in the absence of MNase. Units of MNase (Takara, cat. #2910A) per microgram DNA: lanes 3 and 4, 2.5; lanes 5 and 6, 5.0; lanes 7 and 8, 10, lanes 9 and 10, 20. (F) Agarose gel electrophoreses of di-nucleosomes constructed with c-*fos*-derived annealing sequence DNA. Lane M, DNA ladder marker (NEB, cat. N3232S); lane 1, di-nucleosome reconstituted with unmodified H4; lane 2, di-nucleosome reconstituted with K5/K8/K12/K16-tetra-acetylated H4.

### Reconstitution of di-nucleosome chromatin templates

Di-nucleosomes with or without histone H4 acetylation were reconstituted by salt dialysis (32). Briefly, purified histone octamers were mixed with the X5S 197-2 or X5S 217F-2 fragment DNA in 10 mM Tris-HCl buffer (pH 7.5) containing 2 M NaCl, 1 mM EDTA, and 1 mM 2-mercaptoethanol (histone octamer: DNA = 0.9 : 1.0 w/w), and dialyzed at 4 °C for 16 h. Then, stepwise dialysis was performed against buffers with decreasing NaCl concentrations. The reconstituted di-nucleosomes were fractionated by centrifugation in 10%–25% sucrose gradients at 36,000 rpm at 4 °C for 16 h in a Beckman SW41 rotor. Forty fractions were collected using a Piston Gradient Fractionator (BioComp) and electrophoresed in 0.5× TBE buffer at 8.5 V/cm in a 0.7% Seakem GTG agarose gel and visualized by ethidium bromide staining. Di-nucleosomes from the 20th to 22nd fractions were pooled, concentrated using an Amicon Ultra 0.5 ml 10K centrifugal filter, and dialyzed against 10 mM HEPES buffer (pH 7.5) containing 100 μM EDTA; their concentration was determined from the optical density at a wavelength of 260 nm.

### Atomic force microscopy

Di-nucleosomes (90 ng/μl) were fixed with 0.1% glutaraldehyde for 16 h at 4 °C. Immediately before measurement, they were diluted to 0.45 ng/μl with 10 mM HEPES buffer (pH 7.5), placed onto APTES-treated mica (35) and left for 10 min. Imaging was performed using a high-speed AFM system (Nano Live Vision, RIBM, Tsukuba, Japan) with a carbon nanofiber cantilever (BL-AC10FS, Olympus) at a spring constant of 0.1 N/m in solution phase at 27 °C. Images of a 500 × 375-nm area were obtained at 2 s/frame at a resolution of 192 × 144 pixels. DNA length between two nucleosome core particles was traced manually in the images and quantified using ImageJ software (version 1.45s).

### Nuclease digestion assays

DNA of di-nucleosome (100 ng) containing either tetra-acetylated or unmodified histone H4 was digested for 5 min at 22 °C with micrococcal nuclease (MNase) (0.125 to 2.0 units; Takara, cat. #2910A) in 5.5 mM Tris-HCl buffer (pH 7.6) containing 500 μM CaCl2 and 50 μg/ml BSA. Digestion was terminated by the addition of a 20 mM EDTA solution, containing 0.5% (w/v) SDS and 2 μg of proteinase K (Roche, cat. #3115887). DNA fragments were extracted with phenol/chloroform and analyzed by electrophoresis in a non-denaturing 10% polyacrylamide gel. Restriction endonuclease digestion assay was performed essentially as described (36), in the presence or absence of 0.9 μl of a *Xenopus* oocyte nuclear extract (Supplementary Figure S2). RsaI was serially diluted with a 1x CutSmart buffer (NEB, cat. B7204S).

### Transcription *in vitro* and transcript detection by fluorescence correlation spectroscopy

Transcription was performed essentially as described (37). For calibration, end-initiated RNA (475 nucleotides, nt) was transcribed by T7 RNA polymerase from the X5S 217F-2 template containing the T7 promoter (5’-TAATA CGACT CACTA TAGG-3’). In the FCS assay, each reaction mixture (10 μl) contained purified di-nucleosome or naked DNA template (1 μg), 0.9 μl of a *Xenopus* oocyte nuclear extract, 9.5 mM HEPES (pH 7.4), 100 mM NaCl, 48 mM KCl, 6.7 mM MgCl2, 3.6 mM DTT, 90 μM EDTA, 4.5% (v/v) glycerol, 0.9% (w/v) polyvinylpyrrolidone, 0.06% (w/v) BSA, 500 μM NTPs, 1 μM trichostatin A, 5 units of RNase inhibitor (Toyobo, cat. #SIN-201), and 20 nM c-*fos* antisense probe. The c-*fos* antisense probe was synthesized with a 2’ O-Me RNA backbone, which was labeled with Cy3 at its 5’ end. FCS was conducted on a confocal laser-scanning fluorescence microscope (TCS SP8, Leica) equipped with a single molecule detection unit (PicoQuant) at 23 °C. The reaction mixture (5 μl) was covered with a stainless-steel cylinder (inner diameter, 6.2 mm; length, 5 mm), one end of which was covered with a cover glass. Cy3 fluorescence was excited with a green laser (532 nm, Leica), and emitted photons were captured through an objective lens (63×, HC PL APO CS2 1.20 N.A. water, Leica) with a 570DF30 emission filter (Omega). Each 30-s fluorescence fluctuation measurement was performed with an avalanche photodiode (PicoQuant). Total recording time was about 20 min. Autocorrelation was calculated with the SymPhoTime software (PicoQuant). In each measurement, 131 data points (*i.e.*, fluorescence autocorrelation function; FAF) from 0.01 to 813 ms were obtained; a logarithmic time scale was used. The obtained FAF between 0.01 and 813 ms was approximated with SymPhoTime using the autocorrelation function (equation [1]) with one component:

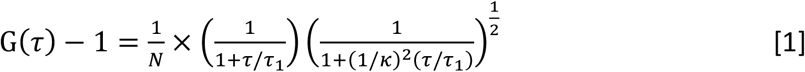

where *N* is the number of fluorescent dyes in the confocal volume, *τ*_1_ is the diffusion time, and *κ* is a structure parameter (10–15 in this experiment).

## RESULTS

### Atomic force microscopy (AFM) imaging and nuclease digestion of the H4-tetra-acetylated di-nucleosome

We reconstituted a di-nucleosome in which two copies of *Xenopus* somatic 5S rRNA gene were connected in tandem (Figure 1A and Supplementary Figure S1) (32) and recombinant human histone H4 was either unmodified or tetra-acetylated at K5/K8/K12/K16 (28,29). Site-specific acetylation of histone H4 at K5/K8/K12/K16 was confirmed by Western blotting using specific monoclonal antibodies (Figure 1B). Histone octamers containing K5/K8/K12/K16-tetra-acetylated H4 (hereafter referred as 4Kac) or unmodified H4 together with bacterially expressed core histones H2A, H2B, and H3 (38) were assembled and purified. Using unmodified or 4Kac histone octamers, 5S rRNA gene di-nucleosomes were reconstituted with either the X5S 197-2 template DNA (Figure 1A, top; Supplementary Figure S1A) (27,32) or the newly designed X5S 217F-2 template DNA (Figure 1A, bottom; Supplementary Figure S1B); X5S 217F-2 had an insertion of a 20-bp human c-*fos*-derived annealing sequence (which is absent in the *Xenopus* genome) at the +115 position downstream of the 5S rRNA gene for specific detection of 5S rRNA nascent transcripts.

Using AFM imaging, we found that the unmodified (Figure 1C) and 4Kac (Figure 1D) X5S 197-2 di-nucleosomes had similar dumbbell-shaped structures. DNA length between the two nucleosome cores did not differ significantly between unmodified di-nucleosomes (41 ± 12 nm; *N* = 50) and 4Kac di-nucleosomes (44 ± 10 nm; *N* = 50). The volumes of the unmodified nucleosome cores (2,100 ± 150 nm^3^; *N* = 10) and 4Kac nucleosome cores (2,000 ± 470 nm^3^; *N* = 10) also did not differ significantly.

We next compared the biochemical accessibilities of micrococcal nuclease (MNase) to the unmodified and 4Kac X5S 197-2 di-nucleosomes (Figure 1E). Toward both di-nucleosomes (424 bp), MNase yielded approximately 145-bp DNA fragments that matched the length of mono-nucleosomal DNA (145–147 bp). The sizes and amounts of this partially digested DNA did not differ between unmodified and 4Kac di-nucleosomes (Figure 1E), suggesting that they had the same chromatin accessibility. Unmodified and 4Kac di-nucleosomes reconstituted with X5S 217F-2 template DNA (Figure 1F) were used for real-time chromatin transcription assay.

### Chromatin transcription and its real-time detection by FCS

To establish a specific system for detection of nascent transcripts by FCS, we first examined naked DNA transcription from the X5S 197-2 and X5S 217F-2 templates in a *Xenopus* oocyte nuclear extract containing RNAP III (Supplementary Figure S3A). We detected 120-nt RNA transcribed from X5S 197-2 and 140-nt RNA transcribed from X5S 217F-2 (Supplementary Figure S3B); thus, transcription from X5S 217F-2 was initiated at the 5S promoter and driven by RNAP III, as described for X5S 197-2 (27). In addition, end-initiated transcripts were also detected from both templates, as reported for X5S 197-2 (27).

To quantify the nascent transcripts, we added an antisense probe (20-nt-long Cy3-labeled 2’-*O*-methyl; 2OMe) (39) to a solution containing 5S rRNA transcripts and the *Xenopus* oocyte nuclear extract in our FCS system; we expected the probe to bind complementarily to the c-*fos*-derived annealing sequence in the transcript (Figure 2A). We assessed hybridization in real time at high resolution as an increase in the diffusion time of the fluorescent antisense probe detected by FCS (40–42) (Figure 2B). However, we could not detect any changes in the diffusion time with the 140-nt RNA, which was transcribed by T7 RNA polymerase, at concentrations ranging from 0 to 10 nM, presumably because the dissociation constant between the 140-nt RNA and the antisense probe is far above 10 nM and/or the difference in sizes between the antisense probe and the hybridized complex was too low to be detected. Therefore, we prepared the end-initiated RNA, which had two copies of 5S rRNA containing the c-*fos*-derived annealing sequence, also by transcription with T7 RNA polymerase. We expected this transcript to increase the sensitivity in FCS owing to its higher affinity to the antisense probe and larger difference in sizes between the antisense probe alone and the probe-hybridized RNA.

**Figure 2.**
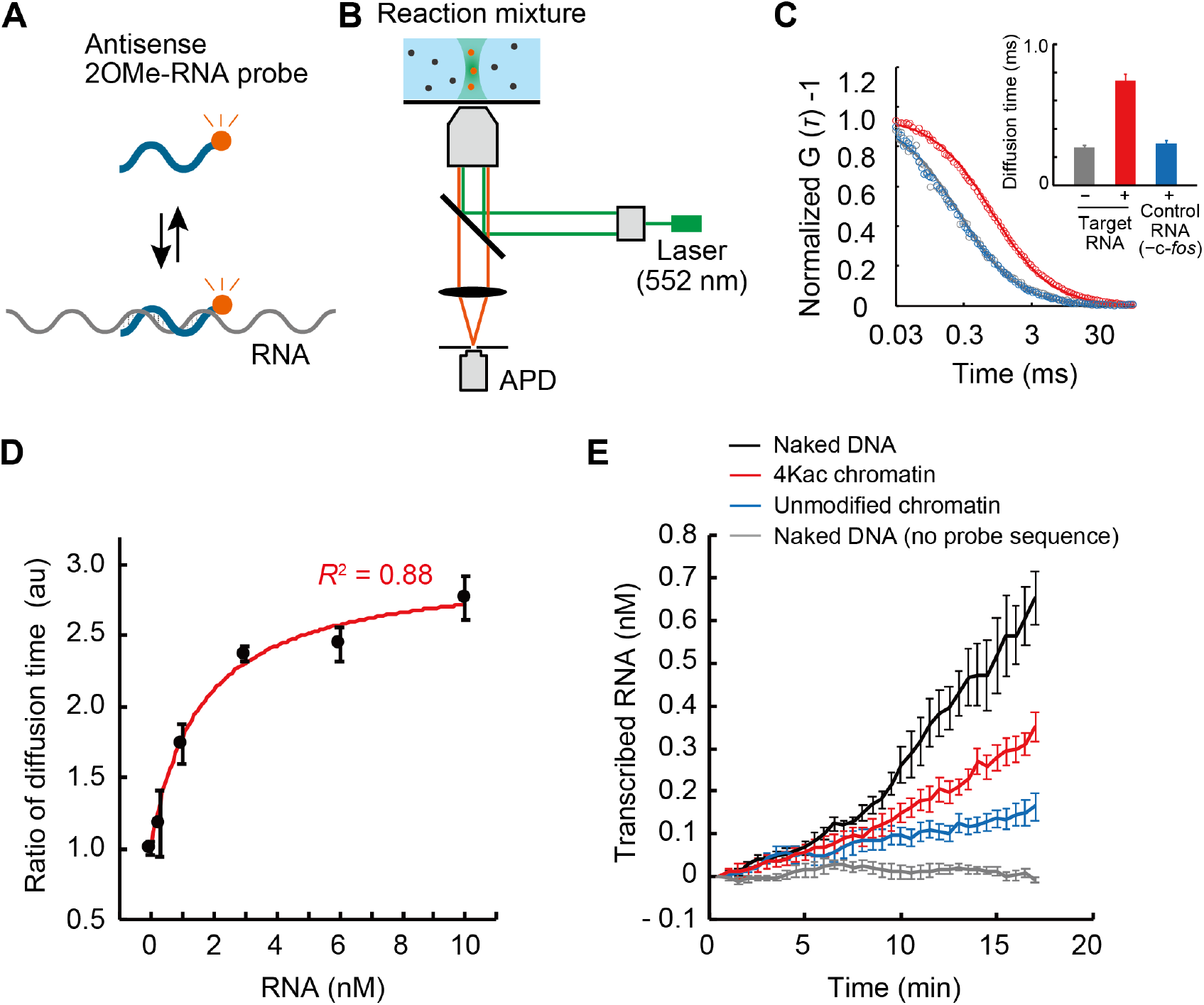
Real-time detection of chromatin transcription by fluorescence correlation spectroscopy. (A) Scheme of a Cy3-labeled antisense 2’-*O*-methyl RNA (2OMe-RNA) probe and its hybridization with mRNA. (B) Scheme of the setup of fluorescence correlation spectroscopy (FCS) for monitoring mRNA synthesis. Changes in diffusion of the antisense probe upon hybridization with transcripts in the confocal volume were detected by an avalanche photodiode (APD) at the single-molecule level. (C) Fluorescence autocorrelation functions [FAF, G(*τ*)] in reaction solutions. FAF of the antisense probe with mRNA showed longer correlation time than that without mRNA or that of a control probe with mRNA. Circles, data points; solid lines, fitting curves. (Inset) Averaged diffusion time of antisense probes. Mean ± standard deviation (*N* = 3). (D) Calibration of the averaged diffusion time of the antisense probe molecules as a function of their concentration. Mean ± standard deviation (*N* = 3). *R^2^,* coefficient of determination. In (C) and (D), end-initiated RNA was transcribed by T7 RNA polymerase from two tandem copies of the 5S rRNA gene (Figure 2A) and added to a *Xenopus* oocyte nuclear extract without template DNA. (E) Real-time detection of nascent 5S rRNA transcripts. Template DNAs used are shown in: gray, naked 5S rRNA gene without c-*fos* sequence; black, naked 5S rRNA gene containing the c-*fos*-derived annealing sequence; red, H4-tetra-acetylated di-nucleosome 5S rRNA gene containing c-*fos* sequence; and blue, non-acetylated di-nucleosome 5S rRNA gene containing c-*fos* sequence. Mean ± standard error (*N* ≥ 3).

When the end-initiated transcript containing two c-*fos*-derived annealing sequences was added to the reaction mixture, the diffusion time of the fluorescent antisense probe, as determined from the FAF, was extended by sequence-dependent hybridization, whereas the end-initiated transcript lacking the c-*fos* sequence did not extend the diffusion time (Figure 2C). To calibrate the transcription level, we added different concentrations of the end-initiated transcript containing two c-*fos*-derived annealing sequences to a reaction mixture that contained all components except the DNA template, incubated the mixture for 10 min, and conducted FCS measurements. The relationship between RNA concentration (0 to 10 nM) and averaged ratio of the diffusion time of the antisense probe to the diffusion time at 0 nM could be approximated by a Michaelis–Menten-type curve with an excellent fit (coefficient of determination, *R*^2^ = 0.88; Figure 2D), confirming that the concentration of the end-initiated nascent transcripts can be quantified by measuring probe diffusion time.

By using a custom-made lid as a cover glass to prevent evaporation, we obtained FAF using as little as 5 μl of the reaction mixtures, thus enabling us to measure the concentrations of nascent transcripts *in vitro* in real time (Figure 2E). We performed transcription assays in the *Xenopus* oocyte nuclear extract with the naked DNA template containing or lacking the c-*fos* sequences, or the X5S 217F-2 di-nucleosome template with tetra-acetylated or non-acetylated H4. As expected, no RNA synthesis was observed for X5S 197-2 naked DNA, which has no probe-annealing sequence, whereas an increase in RNA concentration in a time-dependent manner was observed for X5S 217F-2 naked DNA (Figure 2E). Transcription from the H4-tetra-acetylated di-nucleosome was greater than that from the unmodified di-nucleosome, but both were lower than that from the naked template (Figure 2E). Hence, site-specific acetylation of histone H4 activates transcription from chromatin template *in vitro.*

### Scheme of chromatin transcription and modeling

The reconstituted RNAP III-driven chromatin transcription system used is shown in Figure 3. For analyzing the essential dynamics of chromatin transcription, we postulated four steps on the basis of published data, each representing a chromatin state or reaction: (I) chromatin accessibility, (II) formation of transcriptionally competent chromatin, (III) priming before transcription, and (IV) 5S rRNA transcription.

**Figure 3.**
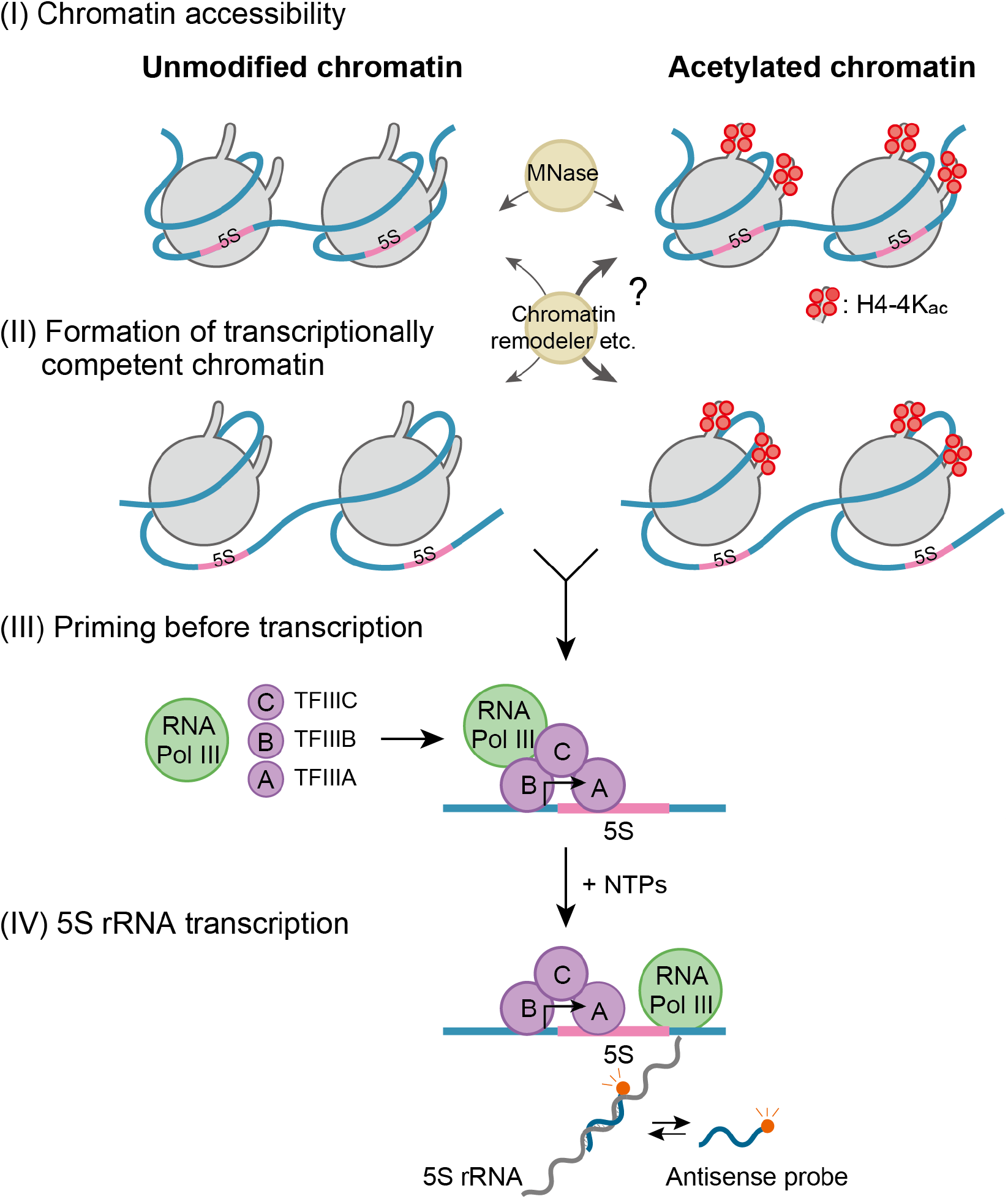
Schematic diagram of 5S rRNA chromatin transcription and its real-time detection.

Chromatin accessibility (I) for *trans*-acting factors can be evaluated by using enzymes that act on DNA, such as micrococcal nuclease (MNase), DNase I, or transposase (43). Chromatin remodeling factors or histone chaperones, or both, may access the chromatin template at different rates, depending on the presence or absence of histone acetylation. Formation of transcriptionally competent chromatin (II) covers a series of possible chromatin-mediated reactions to allow transcription by RNAP III. This step includes reorganization of chromatin around the transcription start site mediated by multi-protein interactions. Priming before transcription (III) includes sequential assembly of the transcription preinitiation complex of general transcription factors TFIIIA/TFIIIB/TFIIIC and RNAP III at the promoter (44–47). Finally, 5S rRNA is transcribed by RNAP III (IV). During RNAP III-dependent transcription elongation, the histone core is moving from a position on the DNA template ahead of RNAP III to a position behind it (48–50). In our model, naked DNA transcription includes steps (III) and (IV) and chromatin transcription includes steps (I)–(IV).

### Kinetic modeling and analyses of chromatin transcription

Using the above chromatin transcription model and FCS data, we constructed a simple kinetic model to quantify: (1) the chromatin transcription kinetics and (2) the contribution of a certain epigenetic modification state (tetra-acetylation of histone H4 in this study) to chromatin transcription. The definitions of the variables and parameters in our kinetic model explained below are summarized in Supplementary Table S1.

At the chromatin accessibility step (I), the concentration of accessible chromatin is expressed by *α_ξ_C*, where *ξ* is the type of chromatin template containing the c-*fos*-derived annealing sequence (‘*ξ*’ = ‘4Kac’ for H4-tetra-acetylated chromatin or ‘unmod’ for unmodified chromatin); *C* (= 300 nM, fixed) is the concentration of the template (‘4Kac’ and ‘unmod’); and *α_ξ_*, is accessibility of chromatin ‘*ξ*’. At the formation of transcriptionally competent chromatin (II) and priming before transcription (III) steps, given *X_ξ_*(*t*) and *Y_ξ_*(*t*) are the concentrations of non-primed transcriptionally competent chromatin ‘*ξ*’ and primed DNA ‘*ξ*’, respectively, at time *t*, the concentration of chromatin ‘*ξ*’ to become transcriptionally competent chromatin is (*α_ξ_C* – *X_ξ_* – *Y_ξ_*) (Figure 4A). The formation of transcriptionally competent chromatin is assumed to be a rate-limiting step and is described as

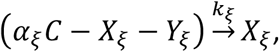

where *k_ξ_* is the rate of transcriptionally competent chromatin formation of chromatin ‘*ξ*’.

**Figure 4.**
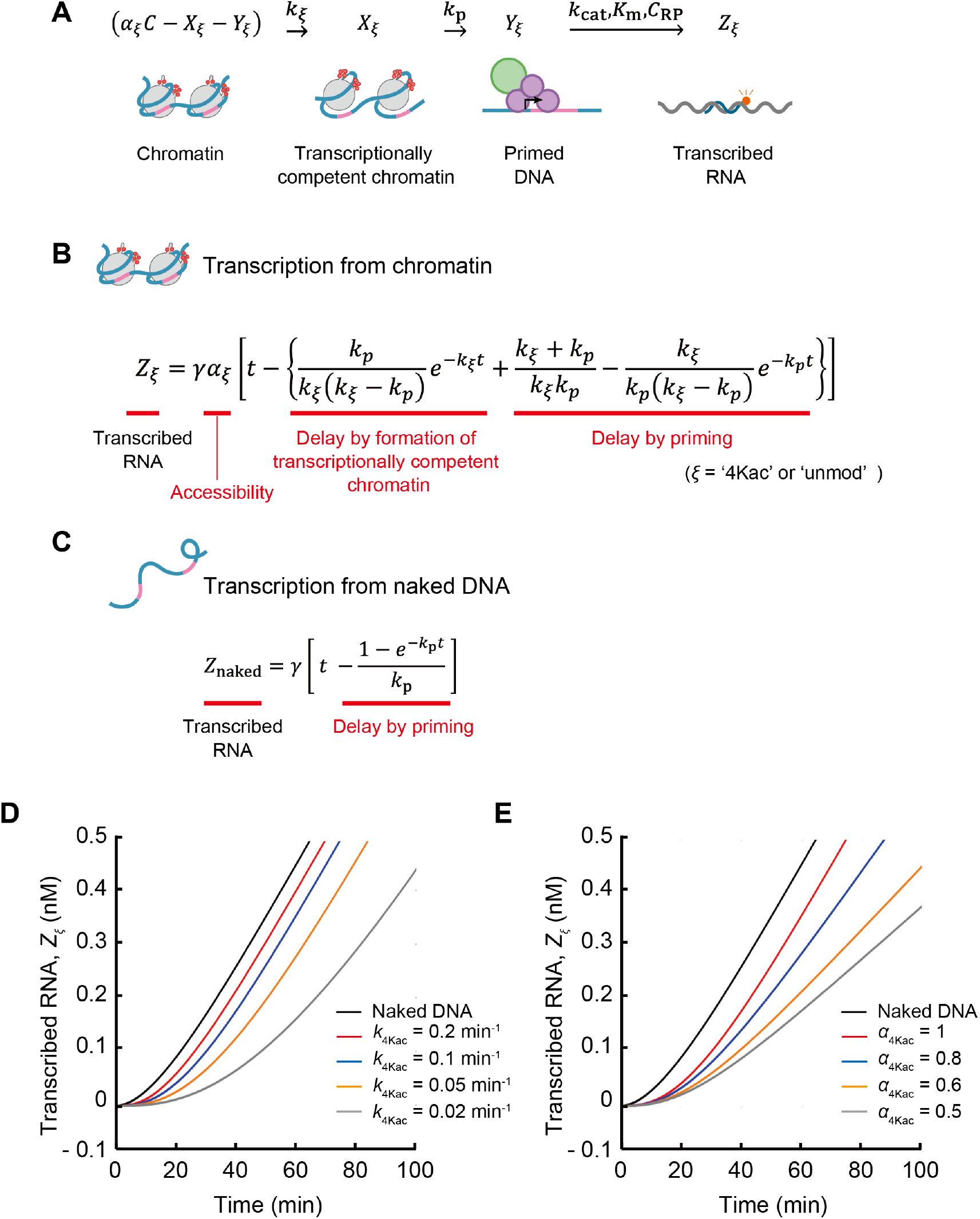
Kinetic model of chromatin transcription. (A) Explanation of modeling of chromatin transcription. Variables are defined in Supplementary Table S1. (B, C) Explanation of the obtained equations. (B) Transcription from chromatin (equation [5]). (C) Transcription from naked DNA (equation [6]). (D, E) Numerical simulation of chromatin transcription with different *k*_4Kac_ (D) and different *α*_4Kac_ (E). *γ* = 0.01 nM min^−1^ and *k_p_* = 1/15 ≈ 0.0667 min^−1^ are fixed. *Z_ξ_* is the concentration of transcribed RNA, where *ξ* is ‘4Kac’ or ‘naked’. For ‘4Kac’, equation [5] is used, where *k*_4Kac_ is changed with *α*_4Kac_ = *α*_naked_ = 1 fixed (D) or *α*_4Kac_ is changed with *α*_naked_ = 1 and *k*_4Kac_ = 0.1 min^−1^ fixed (E). For ‘naked’, equation [6] is used.

Priming before transcription (III) is another rate-limiting step; priming is delayed by sequential assembly of the transcription preinitiation complex at the promoter (16,44,46). Thus, priming before naked DNA transcription is described as

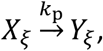

where *k*_p_ is the priming rate of chromatin ‘*ξ*’. Finally, (IV) 5S rRNA transcription reaction is described as follows by assuming the Michaelis–Menten-type enzymatic reaction

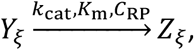

where *Z_ξ_*(*t*) are the concentrations of the transcribed RNA produced from chromatin ‘*ξ*’ at time *t*; *k*_cat_, *K*_m_, and *C*_RP_ are the turnover number, the Michaelis–Menten constant, and the concentration of RNAP III, respectively.

From the chemical reaction model, we have the following ordinary differential equations:

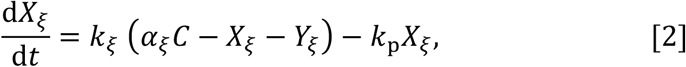

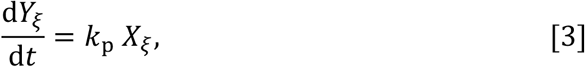

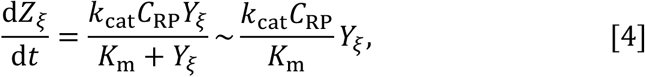

where *K*_m_ ≫ *C* > *Y_ξ_* because *K*_m_ for eukaryotic RNA polymerase III is 7 to 83 μM (51) and *C* in this study was 300 nM. By solving equations [2] – [4] under the initial condition *X_ξ_*(0) = *Y_ξ_*(0) = *Z_ξ_*(0) = 0, the kinetics of 5S rRNA transcription from the H4-tetra-acetylated chromatin and the unmodified chromatin is obtained as

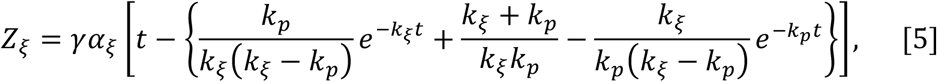

where *ξ* is ‘4Kac’ or ‘unmod’; 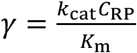 is the naked DNA transcription rate; the term *γα_ξ_t* means pure naked DNA transcription without the need for (II) formation of transcriptionally competent chromatin or (III) priming before transcription; and the term 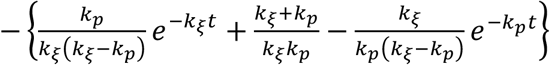 indicates the pre-transcription time delay caused by steps (II) and (III) (Figure 4B).

By taking the limit of *k_ξ_t* → ∞, *k*_p_/*k_ξ_* → 0, and *α_ξ_* = 1 in equation [5], the kinetics of 5S rRNA transcription from the naked DNA containing the c-*fos*-derived annealing sequence are

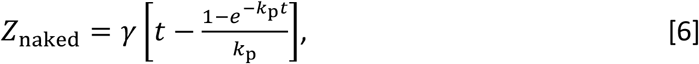

where *Z*_naked_ are the concentrations of the transcribed RNA produced from ‘naked’ DNA at time *t* (Figure 4C).

Using equations [5] and [6], we simulated the dynamics of chromatin transcription involving steps (I) to (IV) with different *k*_4Kac_ (Figure 4D) and different *α*_4Kac_ (Figure 4E). We confirmed that the rate of transcriptionally competent chromatin formation *k_ξ_* contributes to the time delay, not to the final slope of the kinetics (Figure 4D). On the other hand, the accessibility *α_ξ_*, contributes to the final slope of the kinetics, not to the time delay (Figure 4E).

Subsequently, we estimated the kinetic parameters of chromatin transcription by fitting the experimental data to the kinetic model using the computing software Mathematica 11.3 (Wolfram Research, Champaign, IL, USA). First, by fitting the transcription data of the naked DNA to equation [6], we determined the transcription rate *γ* (ca. 0.052 nM min^−1^) and the priming rate *k*_p_ (ca. 0.22 min^−1^; Figure 5A). We used these values to estimate the kinetics of transcription from all types of DNA templates.

**Figure 5.**
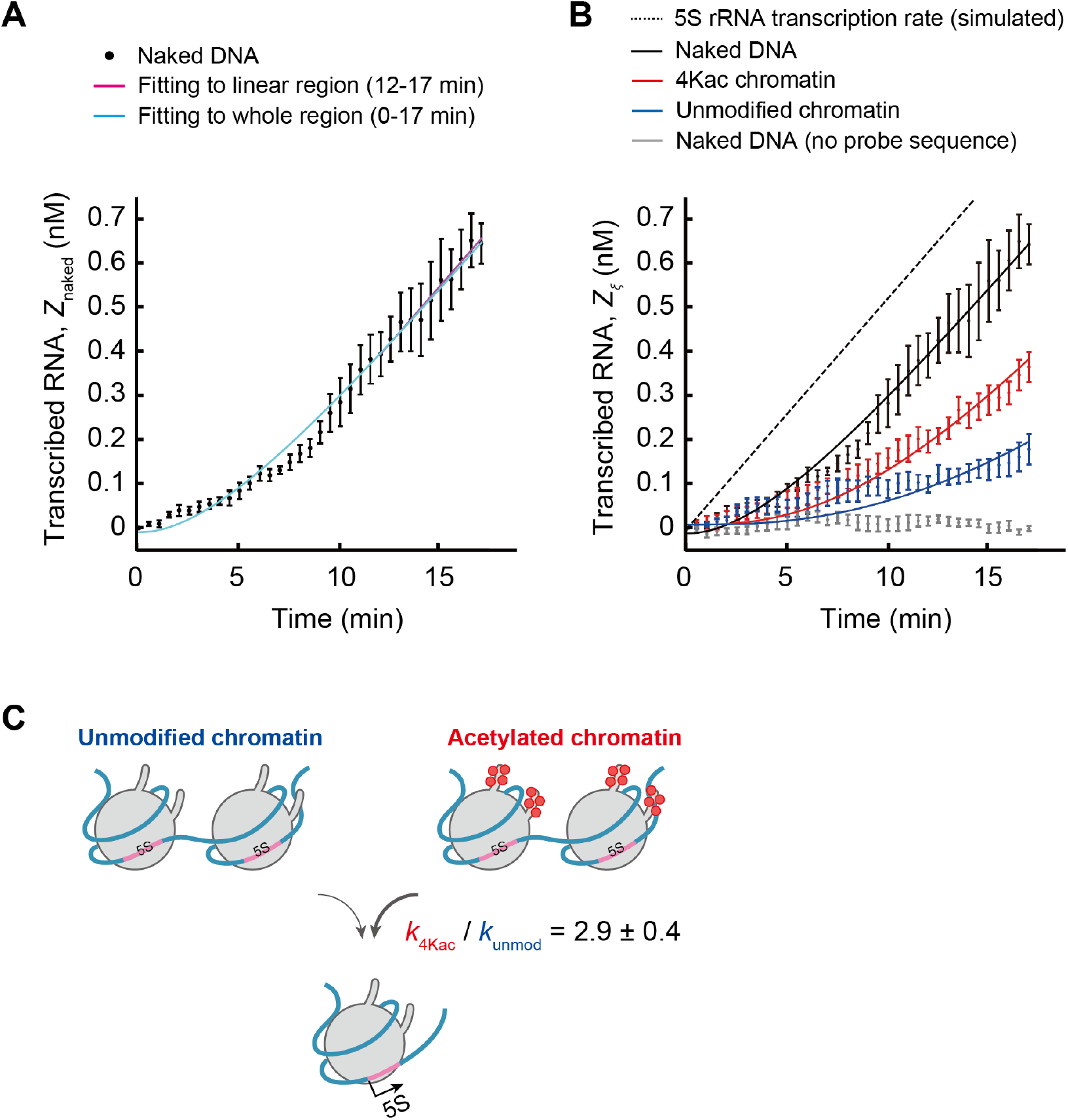
Results of fitting to the kinetic model. (A) Determination of *γ* and *k*_p_. First, the *γ* value was determined by fitting the linear region (12–17 min) of the experimental data for the naked DNA (+ c-*fos*) to the kinetic model *Z*_naked_ = *γt* + *z*_1_, where *z*_1_, is the intercept; this equation is the long-term limit of equation [6]. The fitted values were: *γ* = 0.052 ± 0.003 nM min^−1^ (‘fitting value’ ± ‘fitting error’); *z*_1_ = −0.23 ± 0.05 nM; coefficient of determination *R^2^* = 0.99. Then, using the obtained *γ*, the *k*_p_ value was determined by fitting the whole region (0–17 min) of the data to the kinetic model *Z*_naked_ = *γ*[*t* – (1 – *e*^−*k*_p_*t*^)/*k*_p_] + *z*_2_, where *z*_2_ is the intercept introduced for resolving experimental error at the initial stage (equation [6]). The fitted values were: *k*_p_ = 0.22 ± 0.01 min^−1^; *z*_2_ = −0.010 ± 0.01 nM; *R^2^* = 0.99). (B) Fitting results for each condition. *Z_ξ_* is the concentration of transcribed RNA, where *ξ* is ‘4Kac’, ‘unmod’, or ‘naked’ (equations [5] and [6]). *γ* = 0.052 nM min^−1^ and *k*_p_ = 0.22 min^−1^, which were obtained in (A), were used for fitting. (C) Schematic illustration of acceleration of chromatin transcription by histone H4 acetylation. *k*_4Kac_ ≈ 0.15 ± 0.009 min^−1^ and *k*_unmod_ ≈ 0.052 ± 0.006 min^−1^.

Assuming *α*_4Kac_ = *α*_unmod_ = 1, fitting the 5S rRNA transcription data of the H4-tetra-acetylated and unmodified chromatin to equation [5] (Figure 5B) resulted in the rates of transcriptionally competent chromatin formation *k*_4Kac_ ≈ 0.15 min^−1^ and *k*_unmod_ ≈ 0.052 min^−1^. Because all the coefficients of determination were large (≥0.91), we considered the fitting to have been performed properly (Figure 5A and 5B). This suggests that the kinetic model constructed here can express the transcriptional dynamics. In addition, to examine the adequacy of the assumption of *α*_4Kac_ = *α*_ummod_ = 1, we fitted the data for *α*_unmod_ = 0.8 – 1 (Supplementary Figure S4). Because the *k*_unmod_ and *R^2^* values did not change much even when *α*_ummod_ was changed, the simplest assumption of α_4Kac_ = *α*_unmod_ = 1 was adopted on the basis of the results of MNase digestion assays (Figure 1E). Thus, the mathematically described kinetic analysis shows that H4 tetra-acetylation increases the rate of transcriptionally competent chromatin formation approximately 3-fold in comparison with the absence of modifications (*k*_4Kac_ / *k*_unmod_ = 2.9 ± 0.4; Figure 5C).

## DISCUSSION

Although it has been known since the 1960s that acetylation of N-terminal tails of core histones facilitates chromatin transcription (10,52), the contribution of each histone modification state has not been quantified. In this study, we have developed a methodology for such quantification by a fluorescence-based transcription tracking method in a reconstituted system and kinetic modeling.

To understand the role of a particular epigenetic modification, a protein with this modification has to be produced, for example by enzymatic modification (53), native chemical ligation (54), or genetic code expansion (55,56). To produce H4 histone with four sites specifically acetylated, we combined genetic code expansion and cell-free protein synthesis, but other methods may also be used. By comparing RNAP III-driven transcription dynamics in unmodified and H4-tetra-acetylated di-nucleosomes containing 5S rRNA gene cassettes, we established a platform to evaluate the rates of chromatin transcription involving steps (I) to (IV) (Figure 3).

Around step (II), chromatin remodeling factors such as RSC (remodel the structure of chromatin), or histone chaperones such as FACT (facilitates chromatin transcription), or both, allow access of *trans*-acting factors (*e.g.,* transcriptional machinery) to DNA through relocation of the nucleosome at the transcription start site, which is observed in both RNAP II and III-transcribed genes (57–59). Although we also could not detect significant effects of histone acetylation on chromatin remodeling (Supplementary Figure S2) (20,27,60,61), a certain histone acetylation state might facilitate chromatin reorganization. Furthermore, histone acetylation induces multi-bromodomain-mediated liquid-liquid phase separation (62).

We previously reported that the acetylation of core histones enhances di-nucleosome chromatin transcription by RNAP III without changing nucleosome mobility (27). However, we could not quantitatively analyze the effect of subunit- and site-specific acetylation on chromatin transcription because we prepared histone octamers from HeLa cell chromatin. In the present study, we assessed the site-specific acetylation role of H4 at K5/K8/K12/K16 in chromatin transcription through the genetic code expansion system. Beyond the past study, we intended to quantify: (1) the kinetics of respective step composing chromatin transcription and (2) the contribution of a defined epigenetic modification state to chromatin transcription. Our integrative approach employing genetic code expansion, FCS, and kinetic modeling indicated that the H4 tetra-acetylation increases the rate of transcriptionally competent chromatin formation in 2.9-fold. It would be important to compare the chromatin transcription rates among other H4 acetylation states, such as the newly synthesized K5ac/K12ac species of H4 in the future. The rate may be increased by a combination of H4 acetylation with acetylation of another histone, such as K27-acetylated H3 (15). Because the methodology developed here is versatile, the roles of other histone PTMs (such as methylation and ubiquitination) can also be quantified. The model may also be used for RNAP II-driven chromatin transcription, chromatin replication, and other aspects of DNA metabolism.

Real-time measurement of transcription in crude extracts is challenging because the absolute amount of the reconstituted chromatin template is usually small, which makes conventional measurements in a cuvette difficult. In this study, we overcame this problem by using a fluorescent probe with high affinity and specificity, which was detected by FCS with high sensitivity, and a transcript containing two probe-annealing sequences; this method enables sub-nanomolar real-time measurements of nascent RNA. Measuring molecular diffusion by FCS reduces the likelihood of false positives, which is a problem when solely detecting a change in fluorescence. Furthermore, FCS, based on single-molecule counting, enables detection of RNA newly synthesized from a very small amount of template. We found that minimization of the reaction volume (to <5 μl) and extension of the measurement time (*e.g.,* >20 min) are technical bottlenecks caused by evaporation of the reaction mixture during the measurement, and that fitting data in the present time range (*i.e.,* <20 min) is not easy. Considering the femtoliter order of the FCS measurement region, much smaller volumes and longer measurements can be achieved by integration with microfluidic device technologies. By using the developed transcription tracking method in a reconstituted system, the effects of chemical factors (*e.g.*, other PTMs, ions, pH) and physical factors (*e.g.*, temperature, viscosity, congestion) on the dynamics of chromatin transcription can be investigated. Considering the highly quantitative nature of FCS and the requirement for just a few-microliter reaction volume, the present methodology may also be applicable to a living cell (42).

Our kinetic model quantitatively describes the chromatin transcription steps, although it does not explicitly describe the reorganization of chromatin around the transcription start site, such as chromatin remodeling and nucleosome positioning. We determined the rates of transcriptionally competent chromatin formation *k*_4Kac_ and *k*_unmod_ and confirmed the validity of data fitting, as judged from their biochemically reasonable values. The finding that H4 tetra-acetylation increased the rate of transcriptionally competent chromatin formation approximately 3-fold is important for characterization of the effect of this modification: despite little structural difference from unmodified chromatin, H4-tetra-acetylated chromatin showed considerably more active dynamics of transcription. The effect of the difference observed in this study will be clearer when our methodology is applied to living cells or other PTMs.

The constructed kinetic model (equation [5]) provides insights into transcription from a chromatin template. When focusing on a very short-term process (*k_ξ_t* ≪ 1), 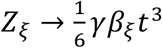, where *β_ξ_* = *α_ξ_k*_p_*k_ξ_* (min^−2^); 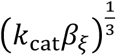 (min^−1^) is an apparent initial rate of transcription from the chromatin template. If very short-term transcription kinetics are quantitatively measured, *β_ξ_* or 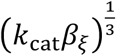 can be used for classifying transcription rates from the chromatin template in various epigenetic modification states. When focusing on a long-term process (*k_ξ_t* ≫ 1,*k*_p_*t* ≫ 1), *Z_ξ_* → *γα_ξ_t*; thus, by measuring the slope of chromatin transcription, chromatin accessibility can be determined more precisely, although in this study it was assumed to be 1 on the basis of the results of MNase digestion assays.

In summary, we established an *in vitro* reconstitution system to quantify the contribution of a certain epigenetic modification state to chromatin transcription dynamics. As a model case, we compared the kinetics of chromatin transcription between unmodified and H4-tetra-acetylated chromatin templates. Chromatin templates with certain PTMs, such as site-specific acetylation, can be used in crude extracts in which the product molecules of interest can be counted in real time. Therefore, our methodology will be applicable to a wide variety of chromatin-mediated reactions for quantitative understanding of the importance of epigenetic modifications.

## SUPPLEMENTARY DATA

Supplementary Data are available at NAR online.

## ACKNOWLEDGEMENTS

We thank Hiroki R. Ueda (the University of Tokyo) for encouragement, Toshiaki Higo (RIKEN) for sample preparation, and Yuki Saito (RIKEN) for clerical assistance.

## Author contributions

M.W., K.O., M.T., and T.U. designed the experiments, interpreted the results, and wrote the manuscript. M.W., K.U., and T.U. performed the biochemical analysis. K.O. and T.F. performed the biophysical analysis. M.T. performed the mathematical analysis. All authors commented on the manuscript.

## REFERENCES

1. Roeder, R.G. and Rutter, W.J. (1969) Multiple forms of DNA-dependent RNA polymerase in eukaryotic organisms. Nature, 224, 234–237.

2. Dignam, J.D., Lebovitz, R.M. and Roeder, R.G. (1983) Accurate transcription initiation by RNA polymerase II in a soluble extract from isolated mammalian nuclei. Nucleic acids research, 11, 1475–1489.

3. Keener, J., Josaitis, C.A., Dodd, J.A. and Nomura, M. (1998) Reconstitution of yeast RNA polymerase I transcription in vitro from purified components. TATA-binding protein is not required for basal transcription. The Journal of biological chemistry, 273, 33795–33802.

4. Vannini, A. and Cramer, P. (2012) Conservation between the RNA polymerase I, II, and III transcription initiation machineries. Molecular cell, 45, 439–446.

5. Paranjape, S.M., Kamakaka, R.T. and Kadonaga, J.T. (1994) Role of chromatin structure in the regulation of transcription by RNA polymerase II. Annual review of biochemistry, 63, 265–297.

6. Felsenfeld, G. (1992) Chromatin as an essential part of the transcriptional mechanism. Nature, 355, 219.

7. Luger, K., Mader, A.W., Richmond, R.K., Sargent, D.F. and Richmond, T.J. (1997) Crystal structure of the nucleosome core particle at 2.8 A resolution. Nature, 389, 251–260.

8. Kornberg, R.D. and Lorch, Y.L. (1999) Twenty-five years of the nucleosome, fundamental particle of the eukaryote chromosome. Cell, 98, 285–294.

9. Peterson, C.L. and Laniel, M.-A. (2004) Histones and histone modifications. Current Biology, 14, R546–R551.

10. Allfrey, V.G., Faulkner, R. and Mirsky, A.E. (1964) Acetylation and Methylation of Histones and Their Possible Role in the Regulation of RNA Synthesis. Proceedings of the National Academy of Sciences of the United States of America, 51, 786–794.

11. Lee, D.Y., Hayes, J.J., Pruss, D. and Wolffe, A.P. (1993) A positive role for histone acetylation in transcription factor access to nucleosomal DNA. Cell, 72, 73–84.

12. Hirschler-Laszkiewicz, I., Cavanaugh, A., Hu, Q., Catania, J., Avantaggiati, M.L. and Rothblum, L.I. (2001) The role of acetylation in rDNA transcription. Nucleic acids research, 29, 4114–4124.

13. Eberharter, A. and Becker, P.B. (2002) Histone acetylation: a switch between repressive and permissive chromatin. Second in review series on chromatin dynamics, 3, 224–229.

14. Grummt, I. and Pikaard, C.S. (2003) Epigenetic silencing of RNA polymerase I transcription. Nature reviews. Molecular cell biology, 4, 641–649.

15. Barski, A., Chepelev, I., Liko, D., Cuddapah, S., Fleming, A.B., Birch, J., Cui, K., White, R.J. and Zhao, K. (2010) Pol II and its associated epigenetic marks are present at Pol III–transcribed noncoding RNA genes. Nature structural & molecular biology, 17, 629.

16. White, R.J. (2011) Transcription by RNA polymerase III: more complex than we thought. Nature reviews. Genetics, 12, 459–463.

17. Wolffe, A.P. (1994) Nucleosome positioning and modification: chromatin structures that potentiate transcription. Trends in biochemical sciences, 19, 240–244.

18. Jiang, C. and Pugh, B.F. (2009) Nucleosome positioning and gene regulation: advances through genomics. Nature Reviews Genetics, 10, 161.

19. Dhalluin, C., Carlson, J.E., Zeng, L., He, C., Aggarwal, A.K. and Zhou, M.M. (1999) Structure and ligand of a histone acetyltransferase bromodomain. Nature, 399, 491–496.

20. Eberharter, A. and Becker, P.B. (2002) Histone acetylation: a switch between repressive and permissive chromatin. Second in review series on chromatin dynamics. EMBO reports, 3, 224–229.

21. Turner, B.M. (1991) Histone acetylation and control of gene expression. Journal of cell science, 99, 13–20.

22. Zhang, K., Williams, K.E., Huang, L., Yau, P., Siino, J.S., Bradbury, E.M., Jones, P.R., Minch, M.J. and Burlingame, A.L. (2002) Histone Acetylation and Deacetylation. Identification of Acetylation and Methylation Sites of HeLa Histone H4 by Mass Spectrometry, 1, 500–508.

23. Smith, C.M., Gafken, P.R., Zhang, Z., Gottschling, D.E., Smith, J.B. and Smith, D.L. (2003) Mass spectrometric quantification of acetylation at specific lysines within the amino-terminal tail of histone H4. Analytical biochemistry, 316, 23–33.

24. Wang, C.I., Alekseyenko, A.A., LeRoy, G., Elia, A.E.H., Gorchakov, A.A., Britton, L.M.P., Elledge, S.J., Kharchenko, P.V., Garcia, B.A. and Kuroda, M.I. (2013) ChIP-mass spectrometry captures protein interactions and modified histones associated with dosage compensation in Drosophila. Nature structural & molecular biology, 20, 202–209.

25. Henry, R.A., Singh, T., Kuo, Y.-M., Biester, A., O’Keefe, A., Lee, S., Andrews, A.J. and O’Reilly, A.M. (2016) Quantitative measurement of histone tail acetylation reveals stage-specific regulation and response to environmental changes during Drosophila development. Biochemistry-Us, 55, 1663–1672.

26. Grunstein, M. (1997) Histone acetylation in chromatin structure and transcription. Nature, 389, 349.

27. Ura, K., Kurumizaka, H., Dimitrov, S., Almouzni, G. and Wolffe, A.P. (1997) Histone acetylation: influence on transcription, nucleosome mobility and positioning, and linker histone-dependent transcriptional repression. The EMBO journal, 16, 2096–2107.

28. Mukai, T., Yanagisawa, T., Ohtake, K., Wakamori, M., Adachi, J., Hino, N., Sato, A., Kobayashi, T., Hayashi, A., Shirouzu, M. et al (2011) Genetic-code evolution for protein synthesis with non-natural amino acids. Biochemical and biophysical research communications, 411, 757–761.

29. Wakamori, M., Fujii, Y., Suka, N., Shirouzu, M., Sakamoto, K., Umehara, T. and Yokoyama, S. (2015) Intra-and inter-nucleosomal interactions of the histone H4 tail revealed with a human nucleosome core particle with genetically-incorporated H4 tetra-acetylation. Scientific reports, 5, 17204.

30. Tse, C., Sera, T., Wolffe, A.P. and Hansen, J.C. (1998) Disruption of higher-order folding by core histone acetylation dramatically enhances transcription of nucleosomal arrays by RNA polymerase III. Mol Cell Biol, 18, 4629–4638.

31. Tanaka, Y., Tawaramoto-Sasanuma, M., Kawaguchi, S., Ohta, T., Yoda, K., Kurumizaka, H. and Yokoyama, S. (2004) Expression and purification of recombinant human histones. Methods, 33, 3–11.

32. Ura, K., Hayes, J.J. and Wolffe, A.P. (1995) A positive role for nucleosome mobility in the transcriptional activity of chromatin templates: restriction by linker histones. The EMBO journal, 14, 3752–3765.

33. Wakamori, M., Umehara, T. and Yokoyama, S. (2012) A tandem insertion vector for large-scale preparation of nucleosomal DNA. Analytical biochemistry, 423, 184–186.

34. Bogenhagen, D.F., Wormington, W.M. and Brown, D.D. (1982) Stable transcription complexes of Xenopus 5S RNA genes: a means to maintain the differentiated state. Cell, 28, 413–421.

35. Wang, H., Bash, R., Yodh, J.G., Hager, G.L., Lohr, D. and Lindsay, S.M. (2002) Glutaraldehyde Modified Mica: A New Surface for Atomic Force Microscopy of Chromatin. Biophysical journal, 83, 3619–3625.

36. Ura, K., Araki, M., Saeki, H., Masutani, C., Ito, T., Iwai, S., Mizukoshi, T., Kaneda, Y. and Hanaoka, F. (2001) ATP - dependent chromatin remodeling facilitates nucleotide excision repair of UV - induced DNA lesions in synthetic dinucleosomes. The EMBO journal, 20, 2004–2014.

37. Birkenmeier, E.H., Brown, D.D. and Jordan, E. (1978) A nuclear extract of Xenopus laevis oocytes that accurately transcribes 5S RNA genes. Cell, 15, 1077–1086.

38. Dyer, P.N., Edayathumangalam, R.S., White, C.L., Bao, Y., Chakravarthy, S., Muthurajan, U.M. and Luger, K. (2004) Reconstitution of nucleosome core particles from recombinant histones and DNA. Methods in enzymology, 375, 23–44.

39. Okabe, K., Harada, Y., Zhang, J., Tadakuma, H., Tani, T. and Funatsu, T. (2011) Real time monitoring of endogenous cytoplasmic mRNA using linear antisense 2’-O-methyl RNA probes in living cells. Nucleic acids research, 39, e20–e20.

40. Kettling, U., Koltermann, A., Schwille, P. and Eigen, M. (1998) Real-time enzyme kinetics monitored by dual-color fluorescence cross-correlation spectroscopy. Proceedings of the National Academy of Sciences, 95, 1416–1420.

41. Medina, M.Á. and Schwille, P. (2002) Fluorescence correlation spectroscopy for the detection and study of single molecules in biology. Bioessays, 24, 758–764.

42. Zhang, J., Okabe, K., Tani, T. and Funatsu, T. (2011) Dynamic association-dissociation and harboring of endogenous mRNAs in stress granules. Journal of cell science, 124, 4087–4095.

43. Tsompana, M. and Buck, M.J. (2014) Chromatin accessibility: a window into the genome. Epigenetics & chromatin, 7, 33.

44. Orioli, A., Pascali, C., Pagano, A., Teichmann, M. and Dieci, G. (2012) RNA polymerase III transcription control elements: Themes and variations. Gene, 493, 185–194.

45. Dieci, G., Fiorino, G., Castelnuovo, M., Teichmann, M. and Pagano, A. (2007) The expanding RNA polymerase III transcriptome. Trends in Genetics, 23, 614–622.

46. Arimbasseri, A.G., Rijal, K. and Maraia, R.J. (2014) Comparative overview of RNA polymerase II and III transcription cycles, with focus on RNA polymerase III termination and reinitiation. Transcription, 5, e27639.

47. Leśniewska, E. and Boguta, M. (2017) Novel layers of RNA polymerase III control affecting tRNA gene transcription in eukaryotes. Open biology, 7, 170001.

48. Clark, D.J. and Felsenfeld, G. (1992) A nucleosome core is transferred out of the path of a transcribing polymerase. Cell, 71, 11–22.

49. Studitsky, V.M., Kassavetis, G.A., Geiduschek, E.P. and Felsenfeld, G. (1997) Mechanism of Transcription Through the Nucleosome by Eukaryotic RNA Polymerase. Science, 278, 1960–1963.

50. Workman, J.L. (2006) Nucleosome displacement in transcription. Genes & development, 20, 2009–2017.

51. Szafranski, P. and Smagowicz, W.J. (1992) Relative affinities of nucleotide substrates for the yeast tRNA gene transcription complex. Zeitschrift fur Naturforschung. C, Journal of biosciences, 47, 320–321.

52. Verdin, E. and Ott, M. (2015) 50 years of protein acetylation: from gene regulation to epigenetics, metabolism and beyond. Nature reviews. Molecular cell biology, 16, 258–264.

53. Mishima, Y., Watanabe, M., Kawakami, T., Jayasinghe, C.D., Otani, J., Kikugawa, Y., Shirakawa, M., Kimura, H., Nishimura, O., Aimoto, S. et al. (2013) Hinge and chromoshadow of HP1alpha participate in recognition of K9 methylated histone H3 in nucleosomes. Journal of molecular biology, 425, 54–70.

54. Fierz, B. and Muir, T.W. (2012) Chromatin as an expansive canvas for chemical biology. Nat Chem Biol, 8, 417–427.

55. Yanagisawa, T., Umehara, T., Sakamoto, K. and Yokoyama, S. (2014) Expanded genetic code technologies for incorporating modified lysine at multiple sites. Chembiochem: a European journal of chemical biology, 15, 2181–2187.

56. Chin, J.W. (2017) Expanding and reprogramming the genetic code. Nature, 550, 53–60.

57. Birch, J.L., Tan, B.C., Panov, K.I., Panova, T.B., Andersen, J.S., Owen-Hughes, T.A., Russell, J., Lee, S.C. and Zomerdijk, J.C. (2009) FACT facilitates chromatin transcription by RNA polymerases I and III. The EMBO journal, 28, 854–865.

58. Helbo, A.S., Lay, F.D., Jones, P.A., Liang, G. and Gronbaek, K. (2017) Nucleosome Positioning and NDR Structure at RNA Polymerase III Promoters. Scientific reports, 7, 41947.

59. Parnell, T.J., Huff, J.T. and Cairns, B.R. (2008) RSC regulates nucleosome positioning at Pol II genes and density at Pol III genes. The EMBO journal, 27, 100–110.

60. Bresnick, E.H., John, S. and Hager, G.L. (1991) Histone hyperacetylation does not alter the positioning or stability of phased nucleosomes on the mouse mammary tumor virus long terminal repeat. Biochemistry-Us, 30, 3490–3497.

61. Klinker, H., Mueller-Planitz, F., Yang, R., Forné, I., Liu, C.-F., Nordenskiöld, L. and Becker, P.B. (2014) ISWI Remodelling of Physiological Chromatin Fibres Acetylated at Lysine 16 of Histone H4. PloS one, 9, e88411.

62. Gibson, B.A., Doolittle, L.K., Schneider, M.W., Jensen, L.E., Gamarra, N., Henry, L., Gerlich, D.W., Redding, S. and Rosen, M.K. (2019) Organization of chromatin by intrinsic and regulated phase separation. Cell, 179, 470–484.e421.

