## Supplementary Data for "Quantification of the effect of site-specific histone acetylation on chromatin transcription rate"

**Supplementary Table S1.** Definition of variables and parameters in the kinetic model.

| Type of input template DNA | Chromatin with tetra-acetylated H4 | Chromatin with unmodified H4 | Naked DNA |
|---|---|---|---|
| Template chromatin concentration | $C$ = 300 nM | | |
| Accessibility for *trans*-acting factors | $\alpha_{4\text{Kac}}$ ($0 < \alpha_{4\text{Kac}} \leq 1$) | $\alpha_{\text{unmod}}$ ($0 < \alpha_{\text{unmod}} \leq 1$) | $\alpha_{\text{naked}} = 1$ |
| Rate of transcriptionally competent chromatin formation | $k_{4\text{Kac}}$ (determined by data fitting) | $K_{\text{unmod}}$ (determined by data fitting) | – |
| Priming rate | $k_{\text{p}}$ (determined by data fitting) | | |
| Transcriptionally competent chromatin concentration | $X_{4\text{Kac}}$ | $X_{\text{unmod}}$ | $X_{\text{naked}}$ |
| Primed DNA concentration | $Y_{4\text{Kac}}$ | $Y_{\text{unmod}}$ | $Y_{\text{naked}}$ |
| Transcribed RNA concentration | $Z_{4\text{Kac}}$ | $Z_{\text{unmod}}$ | $Z_{\text{naked}}$ |
| RNA polymerase concentration | $C_{\text{RP}}$ (nM) | | |
| Turnover number of RNA polymerase | $k_{\text{cat}}$ ($\text{min}^{-1}$) | | |
| Michaelis–Menten constant of RNA polymerase | $K_{\text{m}}$ (nM) | | |
| 5S RNA transcription rate | $\gamma = \frac{k_{\text{cat}} C_{\text{RP}}}{K_{\text{m}}} C$ (nM $\text{min}^{-1}$) (determined by data fitting) | | |

### Supplementary Figure S1

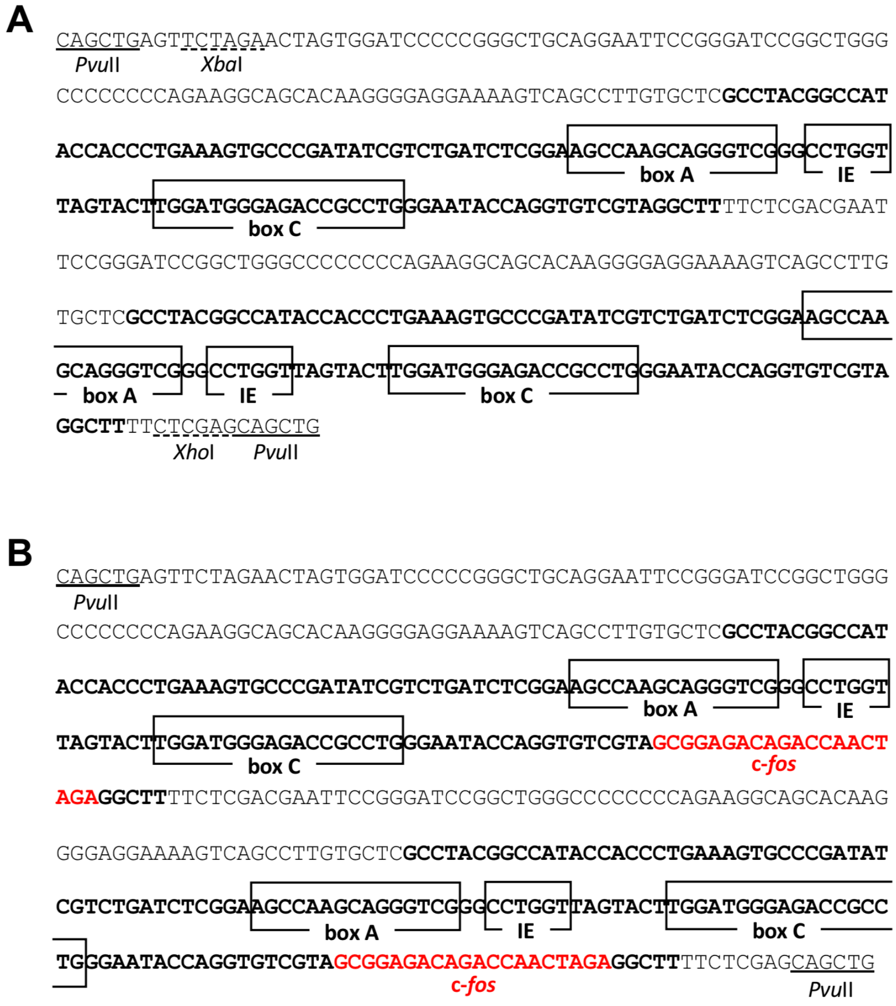


**Supplementary Figure S1.** Chromatin template DNA. (A) Nucleotide sequence of X5S 197-2. (B) Nucleotide sequence of X5S 217F-2. Boxes show internal control regions (box A; IE, intermediate element; box C) of the 5S rRNA gene. A 20-bp human c-*fos*-derived annealing sequence introduced at the +115 and +332 positions downstream of the 5S rRNA gene is shown in red. 5S rRNA gene body is shown in black bold. Restriction enzyme sites (PvuII, XbaI, and XhoI) are underlined.

**Supplementary Figure S2**

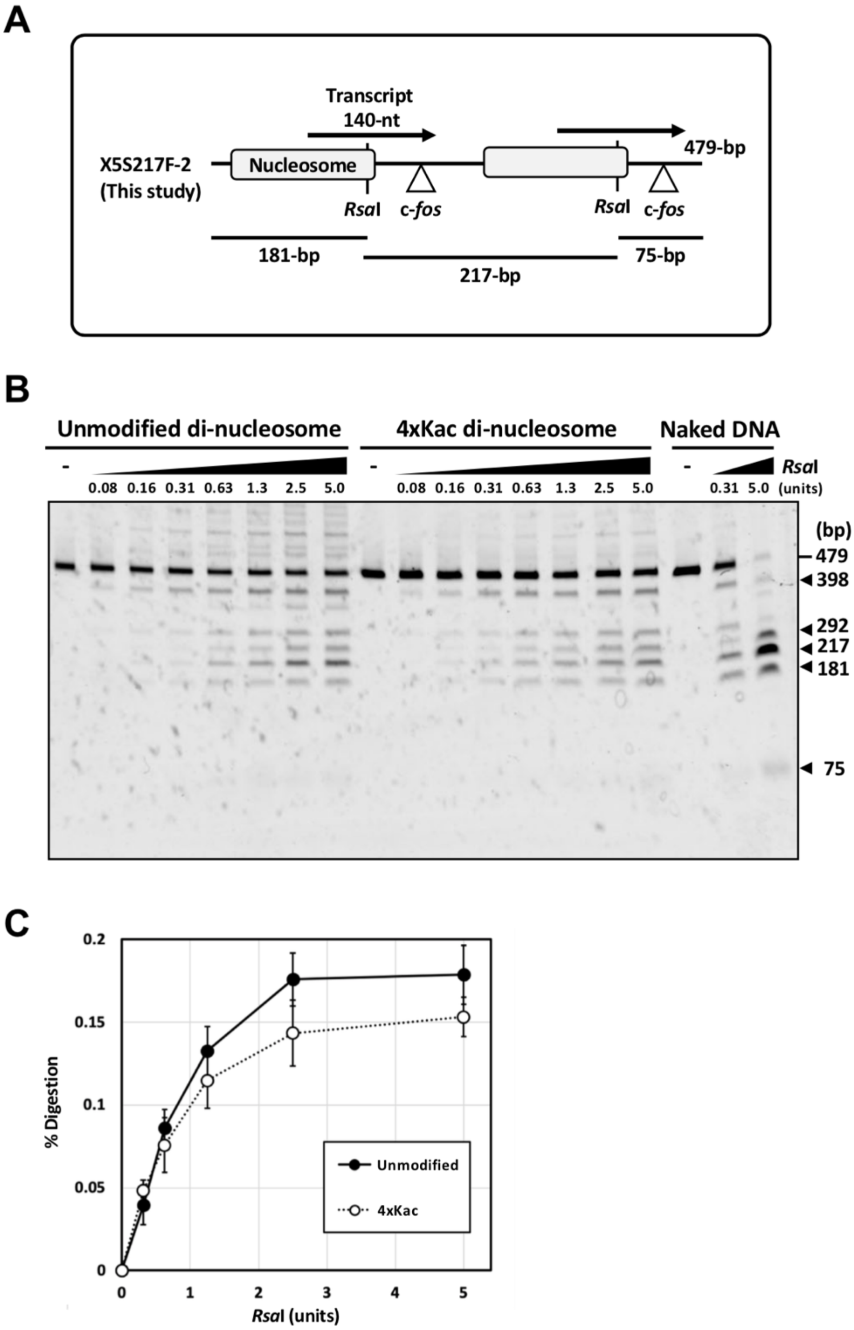


**Supplementary Figure S2.** RsaI digestion assay. (A) Scheme of the digestion. (B) Gel electrophoresis image of the assay. Each reaction mixture (10 µl) contains 50 ng of DNA in the form of unmodified di-nucleosome, H4-tetra-acetylated di-nucleosome, or naked DNA. It also contains 0.9 µl of a *Xenopus* oocyte nuclear extract (NE), 2.5 mM ATP, 10 mM HEPES (pH 7.4), 50 mM KCl, 7 mM $MgCl_2$, 2.5 mM DTT, 100 µM EDTA, 5% (v/v) glycerol, 1% (w/v) polyvinylpyrrolidone, and 10 µM trichostatin A, and was preincubated at 25 °C for 60 min. Then, they were digested for 60 min at 25 °C with RsaI (NEB, cat. #R0167S). The reaction was terminated by the addition of a 20 mM EDTA solution, containing 0.5% (w/v) SDS and 10 µg of proteinase K (Roche, cat. #3115887). DNA fragments were extracted with 25:24:1 phenol-chloroform-isoamyl alcohol (pH 7.9) and analyzed by electrophoresis in a non-denaturing 5% polyacrylamide gel. (C) Digestion efficiency for the DNA fragment that appeared at 217-bp. Mean ± standard deviation ($N$ = 3). $p > 0.05$ at all concentration points.

**Supplementary Figure S3**

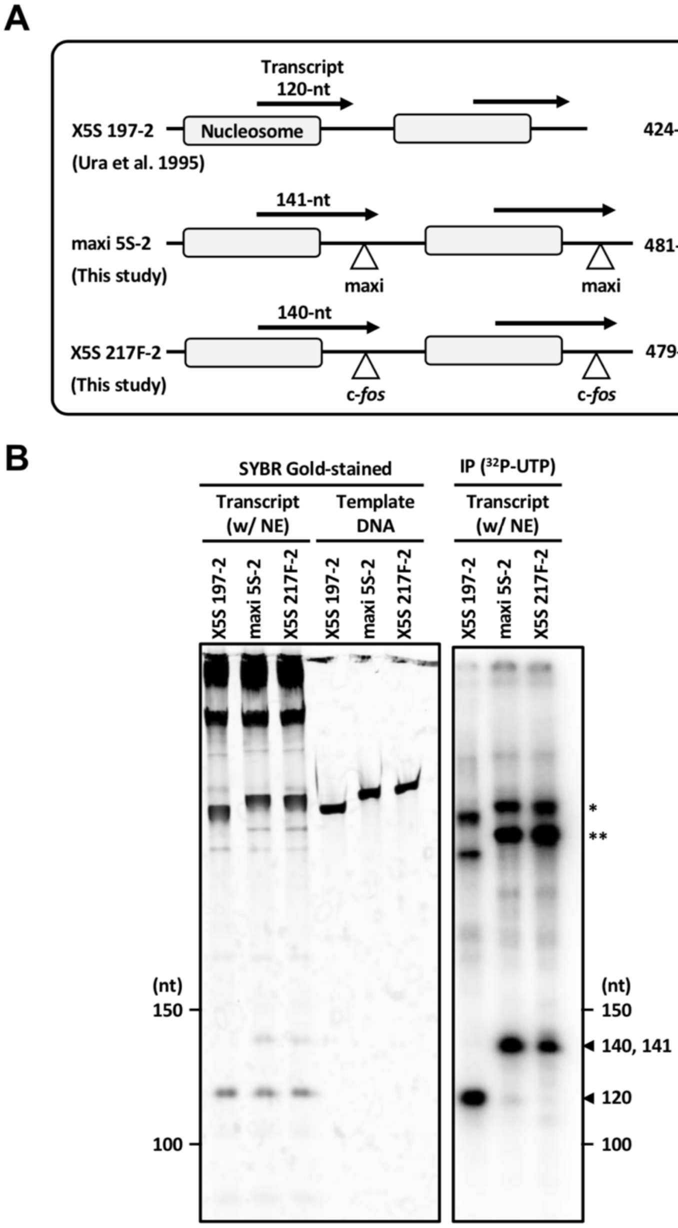


**Supplementary Figure S3.** Naked DNA transcription. (A) Scheme of transcription from X5S 197-2, maxi 5S-2 consisting of two tandem copies of the maxi 5S gene, and X5S 217F-2. The maxi 5S-2 template was constructed for FCS detection using a 21-nucleotide-long annealing sequence, maxi (5'-CCGGA TCCGG CAGGT GTCGT A-3'). (B) Gel electrophoresis images of naked DNA transcription. Each reaction mixture (10 µl), containing naked DNA template (50 ng) , 0.9 µl of a *Xenopus* oocyte NE, 10 mM HEPES (pH 7.4), 50 mM KCl, 7 mM $MgCl_2$, 2.5 mM DTT, 100 µM EDTA, 5% (v/v) glycerol, 1% (w/v) polyvinylpyrrolidone, 1 µM trichostatin A, and 5 units of RNase inhibitor (Toyobo, cat. #SIN-201), was preincubated at 25 °C for 20 min. Then, 500 µM ATP, CTP, and GTP, 100 µM UTP, and 83 pM [α-$^{32}$P] UTP (93 kBq) were added and incubated at 25 °C for 40 min. Radiolabeled transcripts were treated with Proteinase K, extracted with 25:24:1 phenol-chloroform-isoamyl alcohol (pH 5.2), and analyzed by electrophoresis in a 6% denaturing polyacrylamide gel. * Template DNA; ** end-initiated transcripts.

**Supplementary Figure S4**

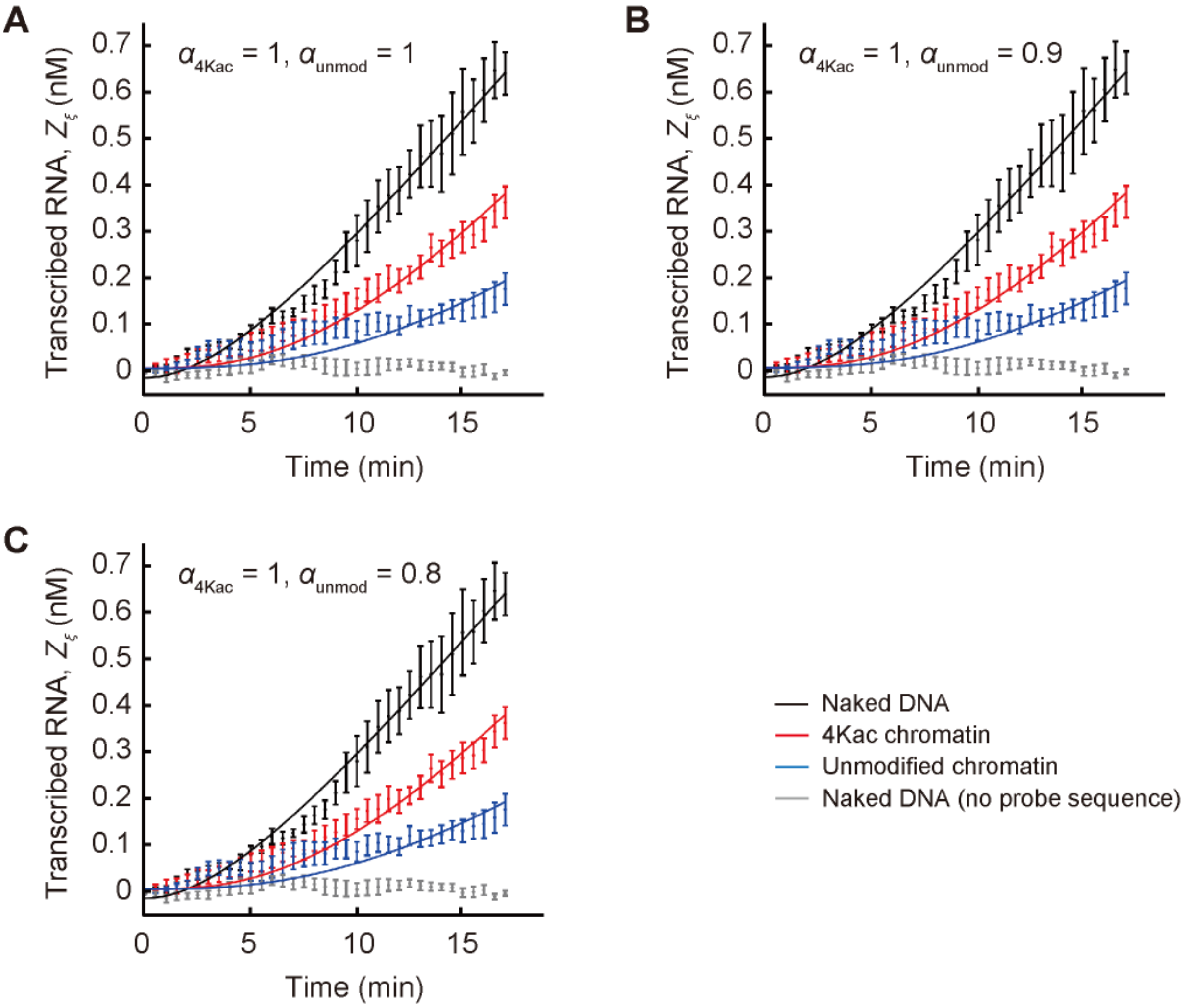


**Supplementary Figure S4.** Data fitting at varying accessibility ($\alpha_{unmod}$). $Z_\xi$ is the concentration of transcribed RNA, where $\xi$ is "4Kac", "unmod", or "naked". For $\xi$ = "4Kac" or "unmod", the equation $Z_\xi = \gamma\, \alpha_\xi \left[ t - \left\{ \frac{k_p}{k_\xi (k_\xi - k_p)} e^{-k_\xi t} + \frac{k_\xi + k_p}{k_\xi k_p} - \frac{k_\xi}{k_p (k_\xi - k_p)} e^{-k_p t} \right\} \right] + z_{3(\xi)}$ was used for fitting, where $z_{3(\xi)}$ is the intercept introduced to resolve experimental error at the initial stage (equation [5] in the main text). The $\gamma$ and $k_p$ values obtained in Figure 5A were used for data fitting. For $\xi$ = "naked", $Z_{naked} = \gamma\left[ t - \left(1 - e^{-k_p t}\right)/k_p \right] + z_2$ was used (equation [6]); $Z_{naked}$ is the replot of Figure 5A. Because $\alpha_{4Kac}$ = 1 was fixed in all panels, we obtained $k_{4Kac}$ = 0.15 ± 0.009 $min^{-1}$ ("fitting value" ± "fitting error"), $z_{3(4Kac)}$ = 0.010 ± 0.007 nM, and coefficient of determination $R^2$ = 0.98 in all cases. (A) $\alpha_{unmod}$ = 1: $k_{unmod}$ = 0.052 ± 0.006 $min^{-1}$ ($z_{3(unmod)}$ = 0.010 ± 0.009 nM; $R^2$ = 0.91). (B) $\alpha_{unmod}$ = 0.9: $k_{unmod}$ = 0.060 ± 0.007 $min^{-1}$ ($z_{3(unmod)}$ = 0.010 ± 0.009 nM; $R^2$ = 0.91). (C) $\alpha_{unmod}$ = 0.8: $k_{unmod}$ = 0.070 ± 0.008 $min^{-1}$ ($z_{3(unmod)}$ = 0.010 ± 0.009 nM; $R^2$ = 0.91). Even if $\alpha_{unmod}$ is changed, $k_{unmod}$ does not change drastically.
